# SEMI-PARAMETRIC COVARIATE-MODULATED LOCAL FALSE DISCOVERY RATE FOR GENOME-WIDE ASSOCIATION STUDIES

**DOI:** 10.1101/183384

**Authors:** Rong W. Zablocki, Richard A. Levine, Andrew J. Schork, Shujing Xu, Yunpeng Wang, Chun C. Fan, Wesley K. Thompson

## Abstract

While genome-wide association studies (GWAS) have discovered thousands of risk loci for heritable disorders, so far even very large meta-analyses have recovered only a fraction of the heritability of most complex traits. Recent work utilizing variance components models has demonstrated that a larger fraction of the heritability of complex phenotypes is captured by the additive effects of SNPs than is evident only in loci surpassing genome-wide significance thresholds, typically set at a Bonferroni-inspired *p* ≤ 5 x 10^-8^. Procedures that control false discovery rate can be more powerful, yet these are still under-powered to detect the majority of non-null effects from GWAS. The current work proposes a novel Bayesian semi-parametric two-group mixture model and develops a Markov Chain Monte Carlo (MCMC) algorithm for a covariate-modulated local false discovery rate (*cmfdr*). The probability of being non-null depends on a set of covariates via a logistic function, and the non-null distribution is approximated as a linear combination of B-spline densities, where the weight of each B-spline density depends on a multinomial function of the covariates. The proposed methods were motivated by work on a large meta-analysis of schizophrenia GWAS performed by the Psychiatric Genetics Consortium (PGC). We show that the new cmfdr model fits the PGC schizophrenia GWAS test statistics well, performing better than our previously proposed parametric gamma model for estimating the non-null density and substantially improving power over usual fdr. Using loci declared significant at cmfdr ≤ 0.20, we perform follow-up pathway analyses using the Kyoto Encyclopedia of Genes and Genomes (KEGG) *homo sapiens* pathways database. We demonstrate that the increased yield from the cmfdr model results in an improved ability to test for pathways associated with schizophrenia compared to using those SNPs selected according to usual fdr.

## 1. Introduction

While genome-wide association studies (GWAS) have discovered thousands of risk loci for heritable disorders, so far even large meta-analyses have recovered only a fraction of the heritability of most complex traits. Recent work utilizing variance components models Purcell et al. (2009); Yang et al. (2010); Davies et al. (2011); Yang et al. (2015) has demonstrated that a much larger fraction of the heritability of complex phenotypes is captured by the additive effects of common variants than is evident only in loci surpassing genome-wide significance thresholds. Thus, the emerging picture is that traits such as these are highly polygenic, and that a large fraction of the heritability is accounted for by numerous loci each with a very small effect (Glazier, Nadeau and Aitman, 2002).

An example is given by the motivating application of this paper, a large meta-analysis of schizophrenia GWAS performed by the Psychiatric Genetics Consortium (PGC, http://www.med.unc.edu/pgc). Schizophrenia is a complex disorder with a heritability (total variability in liability of disease due to variability in genetic factors) estimated from family studies as high as 80%. The latest PGC analyses (Psychiatric-Genomics-Consortium, 2014) combined 82,315 subjects from 52 sub-studies to identify 108 independent regions (128 significant variants) that explained 3% of risk variability. Predictive models using liberally selected collections of thousands of variants not reaching the accepted significance in the PGC study explained as much as 18% of the variability in an independent sample (Psychiatric-Genomics-Consortium, 2014). Further, mixed models used to estimate the total variability in schizophrenia risk explained by all SNP variants tested in the PGC GWAS suggest that as much as 43% of the variability could, in theory, be explained by the collection of variants used for these studies (Psychiatric-GWAS-Consortium, 2011). Taken together these findings suggest that schizophrenia is highly polygenic, with many tiny genetic effects yet to be discovered by conventional statistical approaches and significance criteria, even using more liberal thresholds based on false discovery rate methods (Benjamini and Hochberg, 1995; Efron and Tibshirani, 2002).

Methods for estimating and controlling false discovery rates typically treat all hypothesis tests as exchangeable, ignoring any auxiliary covariates that may influence the distribution of test statistics (Benjamini and Hochberg, 1995; Efron and Tibshirani, 2002). For example, the *local false discovery rate*(fdr) (Efron and Tibshirani, 2002) rests on a simple two-groups mixture model for test statistic *Z*. Letting *f*_0_ and *f*_1_ be the probability density functions corresponding to null and non-null tests, respectively, the marginal pdf of *Z* is given by

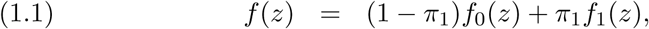

where π_1_ is the non-null proportion. The fdr is then defined as the posterior probability the test is null given the observed test statistic *Z* = *z*.

*Covariate-modulated* fdr (cmfdr) attempts to incorporate the effects of auxiliary covariates into fdr estimation. Ferkingstad et al. (2008) proposed a uniform-beta mixture model for *f*, first stratifying on levels of a scalar covariate *x* and then estimating the parameters of the mixture model within each stratum separately. Lewinger et al. (2007) proposed a noncentral *Χ^2^* distribution for *f*_1_, where the prior proportion π_1_ and the non-centrality parameter are linear combinations of the covariates, passed through nonlinear link functions. Zablocki et al. (2014) proposed a gamma distribution for *f*_1_ where covariates contribute not only to *f*_1_, but also to the prior probability of being non-null. Scott et al. (2015) developed *f*_1_ as a location mixture of null (normal) density and only the prior probability depended on covariates.

These parametric approaches can be efficient if the model fit is adequate. However, the assumed parametric distributions may not always provide an adequate fit to the underlying true non-null distribution, in which case a more flexible nonparametric alternative is desirable to avoid biases in estimating the cmfdr. For example, we found that the gamma distribution underestimated the tails of *f*_1_ in the PGC Schizophrenia GWAS test statistics, leading to elevated estimates of the cmfdr, and hence a loss of power for some loci. The current paper is an extension of Zablocki et al. (2014) to incorporate a more flexible model for the non-null density. We take a semi-parametric approach, modeling the mixture density *f* as a weighted combination of a normal null distribution with B-spline densities bounded away from zero. These non-negative weights are smooth functions of a vector of locus-specific covariates *x*, and normalized to sum to unity. From this mixture model for the density *f*, we can compute a semi-parametric cmfdr, or posterior probability that a test is null given the observed test score *z* and vector of covariates *x*. Model inference is performed via a Markov Chain Monte Carlo (MCMC) sampling algorithm.

Section 2 presents a two-group semi-parametric model for cmfdr incorporating covariates into the estimation of the non-null proportion and density. We describe the MCMC sampling algorithm in Supplementary 1. Section 3 presents Monte Carlo simulations and an application to the PGC Schizophrenia GWAS data. Here, we show large increases in power utilizing functional genomic annotations in the cmfdr model, compared with standard fdr and previous cmfdr methods. The increased yield of SNPs allows for a more powerful pathway analysis of SNPs surpassing a significance threshold of cmfdr ≤0.20. Section 4 concludes with a brief discussion and future directions. The R code for implementing the methods proposed in this paper may be found at https://github.com/rongw16/cmfdr_semi-parametric_model.

## 2. Method

### 2.1. Covariate-modulated local false discovery rate

We use as our starting point the simple two-group mixture model as specified by Eq. (1.1). Let *Z_i_* be random variables, *i* = 1,…,*N*, where *Z_i_* denotes the test statistic for the *i^th^* test. We consider the scenario where for each *Z_i_* we also have an (*M* + 1)-dimensional vector of covariates (including intercept) denoted by *x_i_* = (1, *x*_1*i*_, *x*_2*i*_,…, *x_Mi_*)^*T*^. The test statistics Z_j_ are assumed independent, with marginal density *f* conditional on ***x*** given by

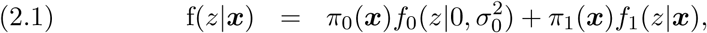

Where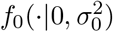 denotes a normal density with mean 0 and variance 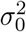 and π_0_(**x**) = 1 — π_1_(*x*). The non-null prior probability *π_1_* and density *f*_1_ depend on the auxiliary covariates *x* as specified in Section 2.2.

We define the cmfdr as the posterior probability that the test is null given Z = *z* and *x*, which by Bayes’ Rule is given by

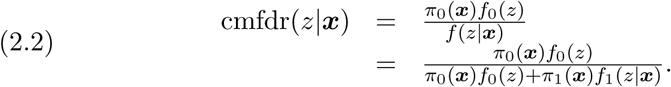

The “zero assumption” of Efron (2007) states that tests with *z*-scores close to zero are primarily of null cases. This is required to ensure the nonnull distribution is identifiable. As in Efron (2007), the default assumption in our applications is that any test with 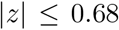 (corresponding to the middle 50% of the standard normal distribution) is considered a null test, i.e., the non-null density *f*_1_(*z*) = 0 for 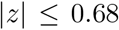. Martin and Tokdar (2012) note that identifiability is not guaranteed for a two-group model with an empirical null involving an unknown variance parameter; however, since a theoretical (standard normal) null poorly describes the behavior of the null in many applications, an empirical null is often required (Scott et al., 2015; Efron, 2004). To solve the problem, Martin and Tokdar (2012) and Scott et al. (2015) impose a “tail assumption” on their models such that *f*_1_ has heavier tails than *f_o_*, where *f_0_* is a normal distribution with unknown mean and variance and *f*_1_ is a location mixture of *f*_0_. We show that our model is identifiable under the zero assumption and other mild conditions (Supplementary 3). In our application of the model to the PGC schizophrenia data, we run multiple chains (each with 23000 iterations) with different random initial values. Figure 1 to 6 in Supplementary 4 depicts convergence of the parameter estimates.

### 2.2. Covariate-modulated mixture density

We first introduce a global latent indicator vector δ = *(δ_1_,…, δ_N_)^T^,* where *δ_i_* = 1 if the *i^th^* test is nonnull and zero otherwise, and *N* is the total number of tests. It is assumed that 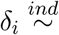 Bernoulli { π_1_ (*x_i_*) }, where

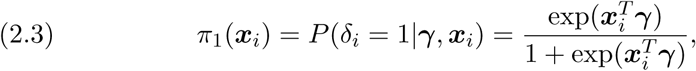

and 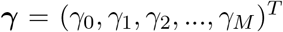 is an (*M* + 1)-vector of unknown parameters. Let *x* denote the (*M + 1) × N* covariate matrix with columns *x_i_*. Then the joint density of *δ* given *γ* and annotations *x* is given by

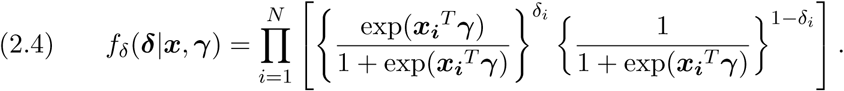

The marginal density of *Z* given by Eq. (2.1) is a mixture of a null density *f*_0_ and a non-null density *f*_1_, each symmetric around zero. Note, the assumption that *f*_0_ and *f*_1_ are symmetric around zero is appropriate for the GWAS example presented here, but could easily be relaxed for other applications. We also assume that *z* scores from null tests are independent and normally distributed with mean zero, that is 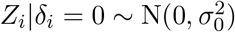. Thus, the

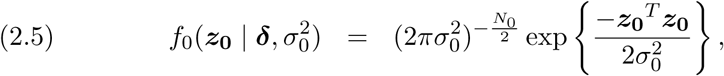

where *N_0_ = N − δ^T^ δ* is the number of tests for which *δ_i_* = 0 and *z_o_* denotes the corresponding *N*_0_-dimensional vector of *z*-scores. The parameter 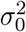 is unknown and estimated from the data (the “empirical null”).

The non-null density *f*_1_ is approximated by a finite mixture of B-spline densities (B-splines normalized to integrate to unity, Lopes and Dias (2012)) with weights that vary smoothly as a function of covariates. B-splines are basis functions having compact support, defined by their polynomial degree and the number and placement of knots (Eilers and Marx, 1996). In the remainder of the paper, we use cubic B-spline densities with knots of multiplicity one fixed by the user, leading to piecewise cubic models with continuous first and second derivatives. Rather than focus on knot selection, the strategy here is to include enough knots to allow a flexible fit and to estimate variance parameters that control the smoothness of the fit (Ruppert, 2002; Thompson and Rosen, 2008).

Specifically, the likelihood of the non-null cases is given by

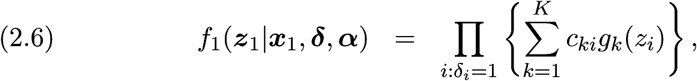

where *z_1_* is the vector of *z*-scores corresponding to non-null tests of dimension; let *N_1_ = δ^T^δ* and *x_1_* is the corresponding *(M + 1) × N_1_* matrix of annotations. The *g_k_* are cubic B-spline densities and the

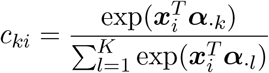

are non-negative weights so that 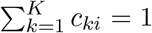. Coefficient *c_ki_* is the probability that the *i^th^* test belongs to the *k^th^* B-spline component, given *δ_i_* = 1 and covariates *x_i_*. These coefficients depend on an (*M + 1) × K* unknown parameter matrix

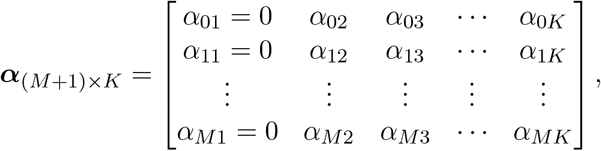

where the *k^th^* column *α._k_* corresponds to the *k^th^* B-spline component and *α_m_*. denotes the row corresponding to the *m^th^* covariate (including intercept), *m = 0,1,2,…, M*. For identifiability, the first column *α._1_*= 0.

We also introduce a local indicator vector 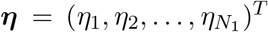. The element 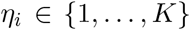 specifies the B-spline component from which the *i^th^* non-null test statistic *z_i_* is generated. The 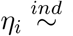 Multinomial(*c_i_*), where *c_i_ = (c_1i_,…, c_Ki_)^T^*. The joint density of *η given δ, α,* and *x_1_* is given by

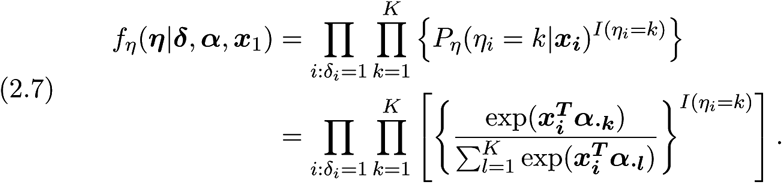

In summary, at the global level, the covariates modulate the probability of the null and non-null status of each test. At the local level (within the non-null distribution), the covariates modulate the B-spline component assignment probability for each non-null test.

#### 2.2.1. Prior distributions

We specify prior distributions for parameters 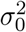, *γ*, and *α*. The rows of *α* are assumed independent. Based on Eilers and Marx (1996), Lang and Brezger (2004), Chib and Jeliazkov (2006), and Rosen and Thompson (2015) we propose the following prior distribution for rows *α_m_*. Let

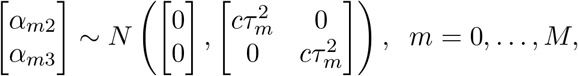

where *c* is a fixed constant and 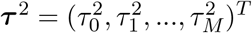 is a (M+1)-vector hyperparameter. In our test runs, *c*=10, 100, or 1000 give similar results; hence *c*=100 is taken in the implementation. The remaining 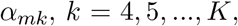, are assumed normally distributed with mean 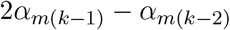 and variance 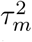 The prior distribution on α_m_. may be expressed in the more compact form as

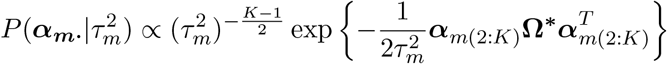

where *α_m(2:K)_* is a (*K* − 1)-vector of B-spline components for the *m^th^* co-variate and *Ω^*^* is a *(K − 1) × (K — 1)* matrix defined as follows. Let

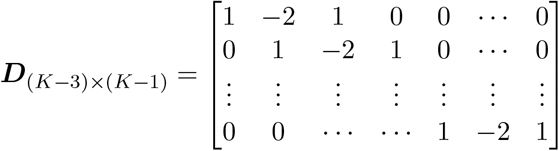

and Ω = *D^T^ D.* We define Ω^*^ = Ω, except for 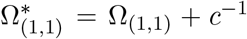 and 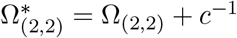 to ensure that the matrix Ω^*^ is positive definite

We propose Inverse Gamma prior for each 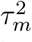 based on Wand et al. (2011), Gelman et al. (2006) and Rosen and Thompson (2015),

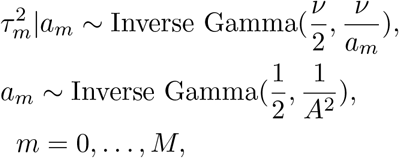

where *a = (a_0_,a_1_,…,a_M_)^T^* is a (M+1)-vector hyperparameter and *a_m_* follows an Inverse Gamma distribution. Hyper-parameters *v* and *A* are assumed known; in our experience, values of *v*, 10 or 20, and values of *A*, 10 or 10,000, yield similar results, as observed in Rosen and Thompson (2015). Therefore, we take *v* = 10 and *A* =10 in the implementation. The kernel probability functions of 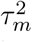 and *a_m_* take the following forms:

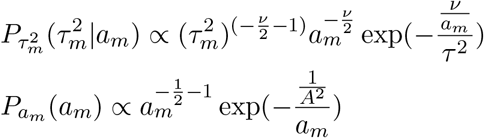

To complete the model, we assume weakly informative priors on the unknown parameters γ and 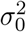:

- 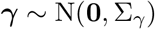
- 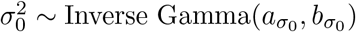

where hyperparameters Σ_#x03B3;_, *a_σ0_*, and *b_σ0_* are fixed by the user. In the simulations and data application, we set Σ_γ_ to be diagonal with variance 10,000 and *(a_σ0_, b_σ0_)* = (0.001, 0.001). Conditional posterior distributions and the MCMC sampling algorithm are described in Supplementary 1.

## 3. Results

### 3.1. Simulation study

In these simulation studies, we set the minimum non-null 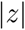-score at 1.96, 0.68, and 0.25 to represent high, medium, and low power scenarios (corresponding to the central 95th, 50th, and 20th percentiles of a standard normal distribution, respectively). We set γ = (−5.29, 2.5, −1.5)*^T^*, γ = (−3.74, 1.2, −1)*^T^*, and ³ = (−3.06, 0.5, −0.2)*^T^* to represent large, medium, and small effects, respectively. These choices for *γ_0_* set the true non-null proportion in all simulations around 5%. The variance parameter 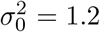 = 1.2. The values for α, *τ^2^* and *a* are drawn from their respective distributions as described in Section 2.2.1.

Each of the nine combinations of power scenarios and covariate effects includes 100 datasets, each dataset includes N=50,000 hypothesis tests where K = 5. Two covariates are generated, with *x*_1_ binomial and *x*_2_ standard normal random variables. We compare the proposed cmfdr model to an intercept only model, which is functionally equivalent to the fdr given in Efron (2007). For each setting each dataset, the MCMC algorithm was run for 18,000 iterations with 1,400 retained samples.

Table 1 presents the median values of sensitivity, specificity, false discovery proportion (FDP, defined as the proportion of incorrectly identified non-null nodes) and number of the non-null cases identified, as well as corresponding 95% credible intervals from 100 runs. Significance cutoffs for both fdr and cmfdr are set to 0.05. Specificity is consistently high and FDP is consistently low across all conditions. Sensitivity and the number of identified non-null cases are consistently higher in cmfdr comparing with fdr (horizontal comparisons) across all conditions. Increased sensitivity is more pronounced with low and medium power regardless of covariate effects. For example, at high power large covariate effect scenario, sensitivity increases 6.9% and 195 more non-null cases are identified by cmfdr comparing to fdr; where as for the medium power/large covariate effect scenario, sensitivity is increased by 14.4% and 400 more non-null cases are identified by cmfdr. These results suggest that in the high power scenario, the null and nonnull distributions tend to be naturally separated, the covariate effects may become less important.

**Table 1.**
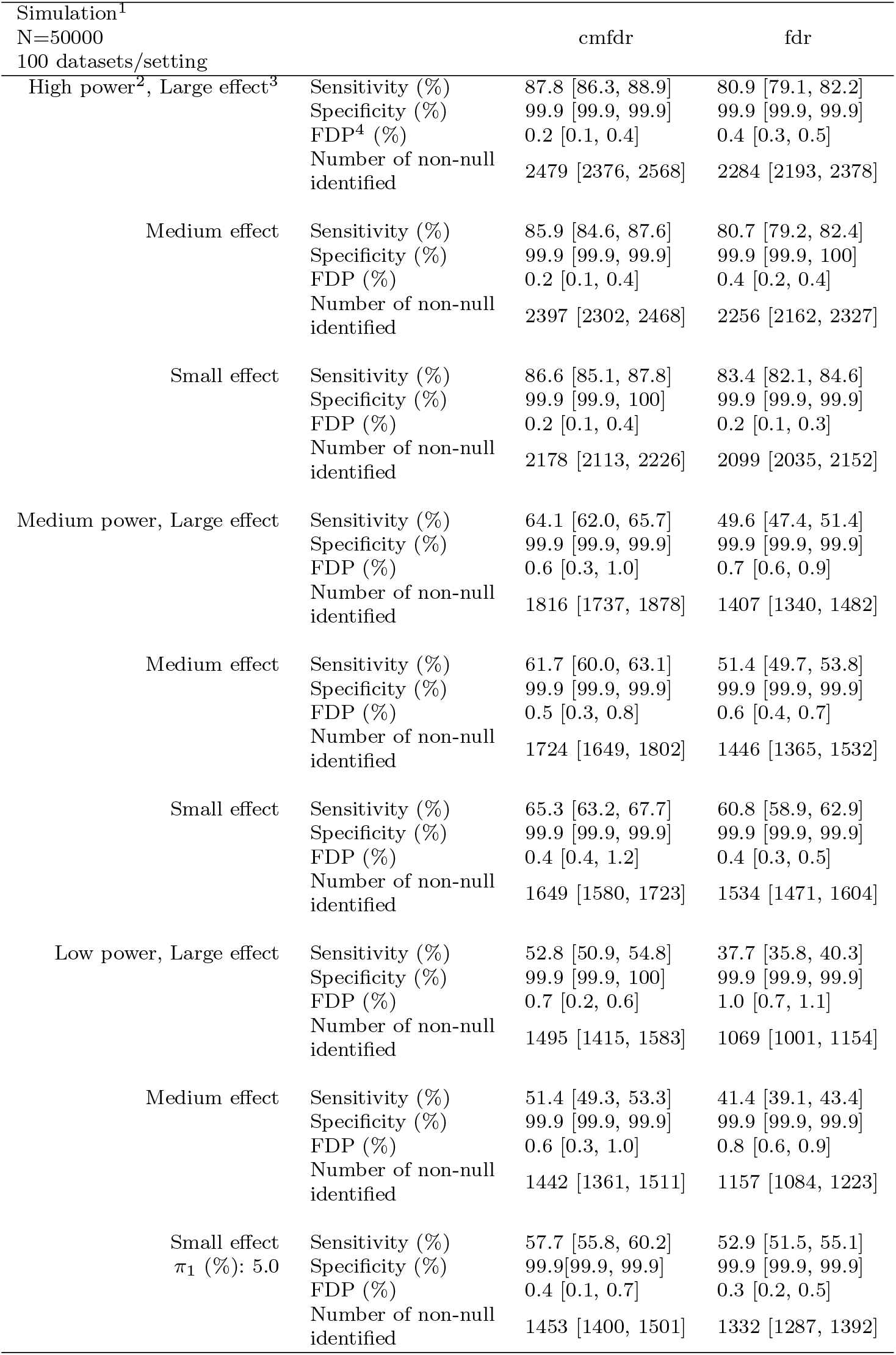
Performance comparison between semi-parametric cmfdr and fdr under different conditions.

### 3.2. Schizophrenia GWAS Application

For this study we used publicly available (https://www.med.unc.edu/pgc/downloads) results from the PGC Schizophrenia GWAS meta-analysis (Psychiatric-Genomics-Consortium, 2014). These data consist of summary statistics for 9,279,485 SNP variants. For each SNP variant independently, a fixed effects meta-analysis was performed across the results of 52 sub-studies. Each sub-study used a logistic regression to test the count of one of the two variant alleles (0, 1 or 2) for association with schizophrenia (as a case-control outcome), adjusted for nuisance co-variates.

The allele counts of variants in close proximity on the genome are correlated (termed Linkage Disequilibrium or LD, Reich et al. (2001)) with the dependence falling off approximately exponentially with distance, although at variable rates across the genome. As a result, the test statistics from a GWAS are not independent and have a variable width, approximately block diagonal correlation structure. To obtain an approximately independent subset of test statistics, we compute the pairwise squared correlation coefficient (*r*^2^) between allele counts for all pairs of SNPs within a conservatively large window of 1,000,000 base pairs. Genotype data for the PGC study were not available, so correlations were estimated in an independent, but representative, collection of European individuals sequenced as part of the 1000 genomes project (Genomes-Project-Consortium, 2012). To facilitate follow-up pathway analyses, we assigned SNPs with gene annotations corresponding to genes within 50,000 base pairs for that given SNP. Genes were selected based on the 242 Kyoto Encyclopedia of Genes and Genomes (KEGG) *homo sapiens* pathways (Kanehisa and Goto, 2000; Kanehisa et al., 2016) SNPs within the major histocompatibility complex (MHC) on chromosome 6 were removed due to the extensive and complex correlation structure within the region. The resulting test statistics were then randomly pruned for approximate independence, such that the estimated squared correlation coefficient *r^2^* was less than 0.2 for any pair of SNPs. In order to approximate the maximum independent set of those SNPs (to minimize the information loss due to the pruning) our pruning scheme is based on a greedy algorithm which in each step keeps a node with the minimum number of neighbors in a complete graph. The final data are composed of *N* = 74,800 SNP summary statistics (*z*-scores) on *n* = 82,315 subjects (35,476 cases). The meta-analysis *z*-scores of the 52 sub-studies are calculated based Willer, Li and Abecasis (2010) and converted to *z*-test statistics using the inverse (standard normal) probability transform.

For each SNP, we also computed three covariates: (1) the Total LD score (TotLD), which is the sum of the squared correlation coefficients between a given SNP and all others within a 1,000,000 base pairs window, again computed in the representative 1000 genomes sample, a measure of the size of the correlation block the SNP resides in; (2) heterozygosity (H), which is the variance of the allele count, or *H = 2(p)(1 — p)*, where *p* is the frequency of the reference allele; (3) the Total Protein Coding Gene LD score (Protein-Coding), which sums the squared correlation coefficients between a given SNP and all others within a 1,000,000 base pairs window that are in a protein coding gene as annotated on the reference genome (Hsu et al., 2006), a rough measure of the functional DNA within a SNP’s correlation block. We have previously shown that these three covariates enrich for non-null SNP associations across a broad range of complex phenotypes (Schork et al., 2013). The distributions of TotLD and ProteinCoding are highly skewed and thus were log-transformed. All three covariates were then standardized to have mean zero and standard deviation one. The MCMC algorithm was applied with K=5. Parameter estimates for γ indicate that all three covariates are positively associated with the prior probability of non-null status in semi-parametric cmfdr, where coefficient for TotLD is 0.73, 95% credible interval is [0.61, 0.86]; H: 0.31 [0.24, 0.38] and ProtenCoding: 0.29 [0.22, 0.37]. The positive association are also observed in gamma cmfdr (Zablocki et al., 2014) as well as in FDRreg cmfdr (Scott et al., 2015).

Power to detect non-null SNPs in different models is displayed in Figure 1. This figure compares the number of non-null SNPs rejected under different models as a function of significance threshold. The increase in power for both the gamma and semi-parametric cmfdr approaches compared to fdr, across a range of cut-offs from 0.001 to 0.20, is large. For example, for cut-off 0.20, fdr rejects 175 null hypotheses, semi-parametric cmfdr with all three covariates rejects 588, gamma model cmfdr rejects 368, and FDRreg cmfdr rejects 203. For reference, the commonly-used GWAS threshold of *p ≤ 5 × 10^−8^* rejects

**Figure 1.**
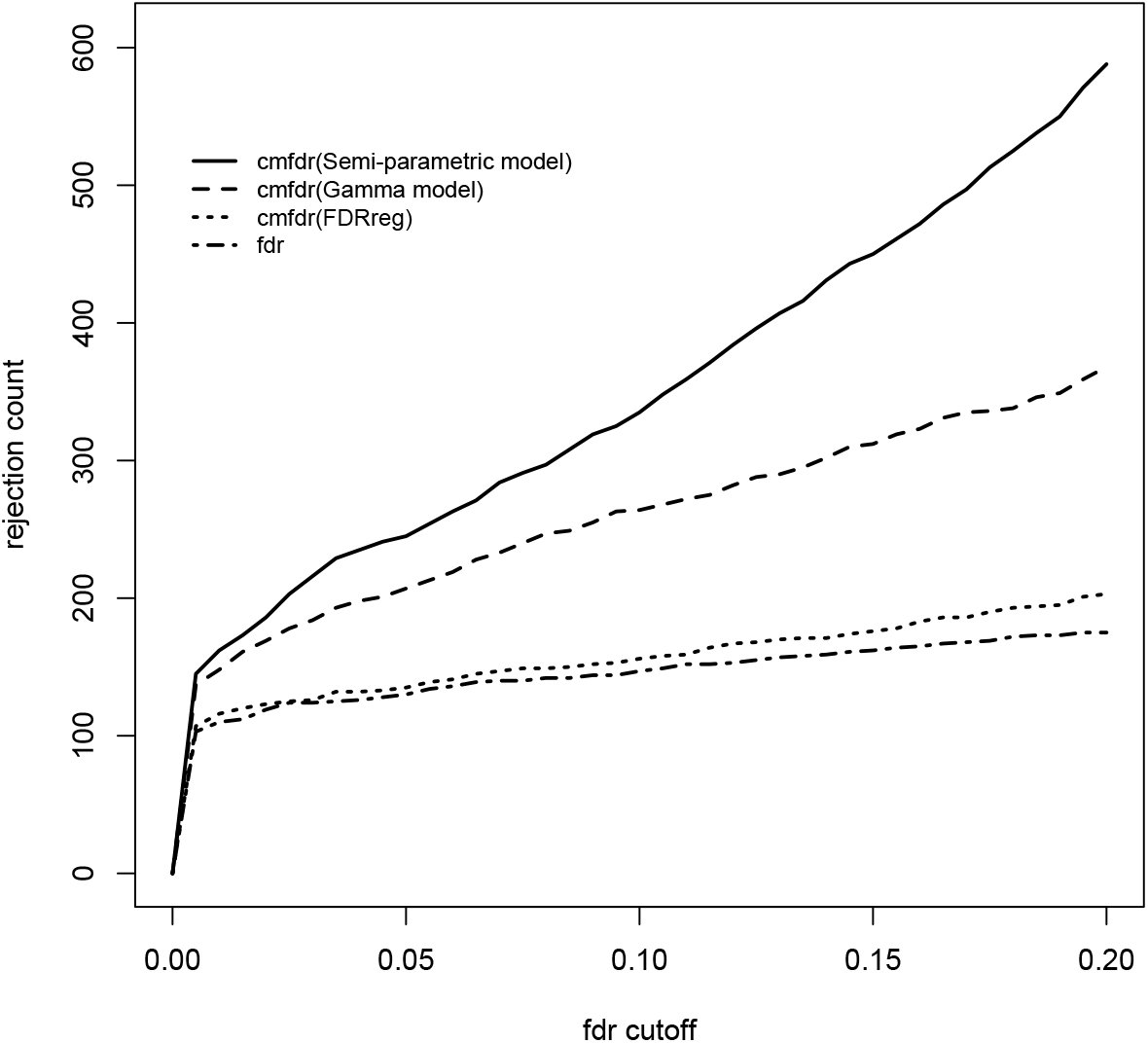
Power curve for fdr, cmfdr (FDRreg), cmfdr (gamma model) and cmfdr (semi-parametric model). The *x*-axis is the fdr cutoff required to declare a SNP significant. The *y*-axis is the number of rejected SNPs.

We also investigate the model fits, comparing semi-parametric cmfdr and the parametric gamma cmfdr. Figure 2 presents stratified Q-Q plots by π_0_ quant.iles. This figure displays the — log_10_ observed *p*-values vs. the theoretical — log_10_ *p*-values under a standard normal distribution. Each SNP has been assigned to one of three strata based on π_0_(***x_i_***) - 1— π^1^(***x_i_*** value by quant.iles: [0.00, 0.33], (0.33, 0.66], and (0.66, 1.00]. The predicted — log_10_ *p*-values estimated from the models are shown with a solid line, dashed line, and dotted line, respectively; the observed — log_10_ *p*-values are shown with dots, triangles, and stars. SNPs in the stratum π_0_ : [0 — 33] have the highest likelihood of being non-null, while SNPs in the stratum πo : (66 — 100] have the highest probability of being null. The gray dash-dot line indicates where the Q-Q curve would lie if all SNPs were null under a standard normal distribution. The leftward deflection of the — log_10_ *p*-values on the Q-Q plots stratified by π_0_ quantiles implies an abundance of non-null SNPs versus the global null hypothesis. The semi-parametric cmfdr displays the best model fit compared to the data. Of the 588 SNPs rejected by the semi-parametric cmfdr at the 0.2 cutoff, 578 are from the stratum π_0_ : [0 — 33], 9 from the stratum π_0_ : (33 — 66], and only one from the stratum π_0_ : (66 — 100]. Analogously, of the 368 SNPs rejected by the gamma cmfdr at the 0.2 cutoff, the numbers of SNPs in corresponding strata are 364, 4, and 0, respectively.

Furthermore, we plot the semi-parametric cmfdr (Figure 3a) and gamma cmfdr (Figure 3b) versus the observed absolute *z*-scores stratified by quantiles of π_0_(*x_i_*); fdr is also added as a reference. The gray dotted line is the 0.2 cutoff. For the most enriched sample, the minimum absolute *z*-scores with semi-parametric cmfdr ≤ 0.2 is 2.25 and with gamma cmfdr ≤ 0.2 is 2.57. For fdr, the minimum absolute *z*-score under this threshold is 4.46, further demonstrating the increase in power from using cmfdr vs. fdr.

**Figure 2.**
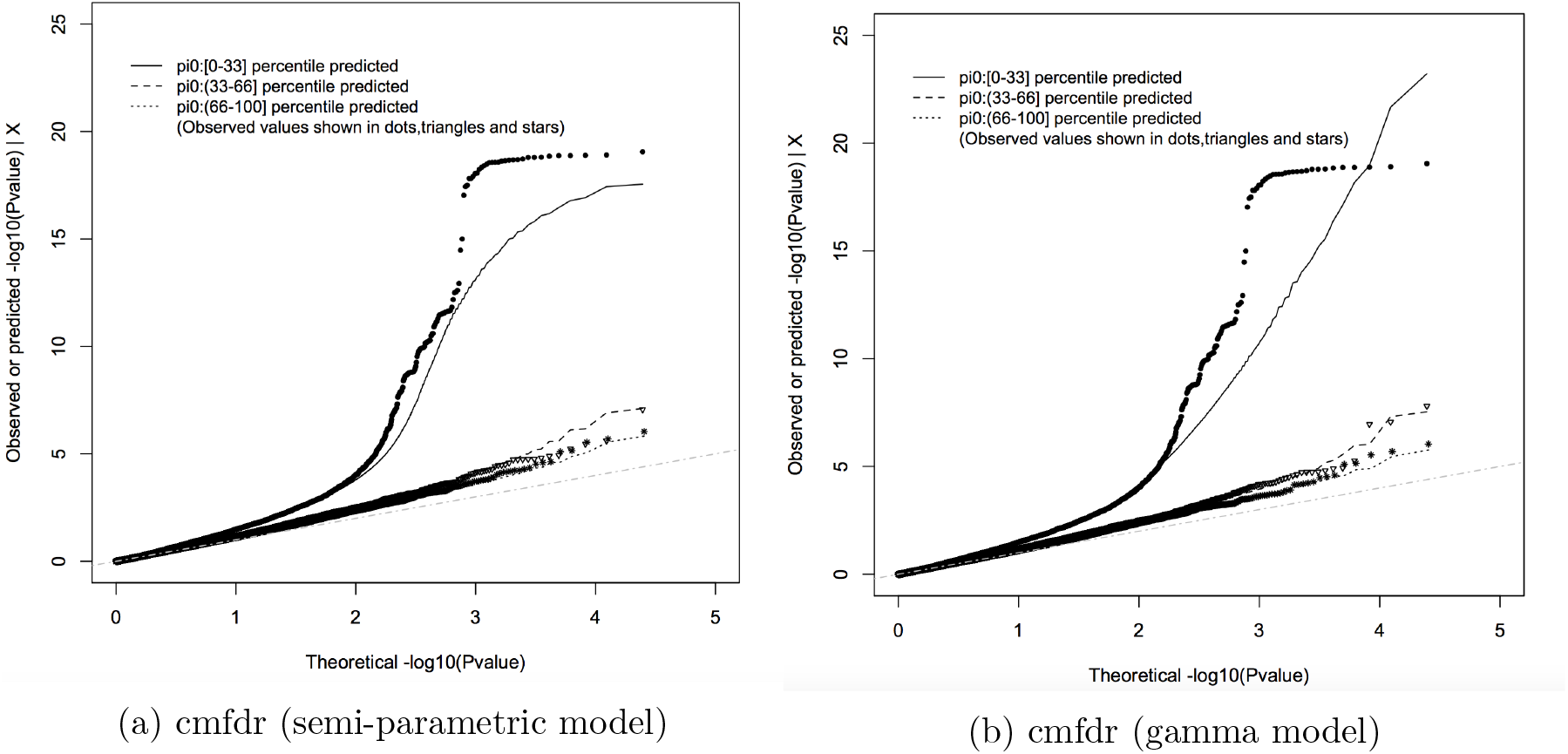
Q-Q plot by π_0_ quantile for the PGC Schizophrenia GWAS data. The *x*-axis is the theoretical −log_10_ *p*-values under a standard normal distribution. The *y*-axis is the −log_10_ observed or predicted *p*-value (converted from *z*-scores). The gray dash-dot line is the reference line indicating where the −log_10_ *p*-values would lie if all SNPs were null under a standard normal distribution.

**Figure 3.**
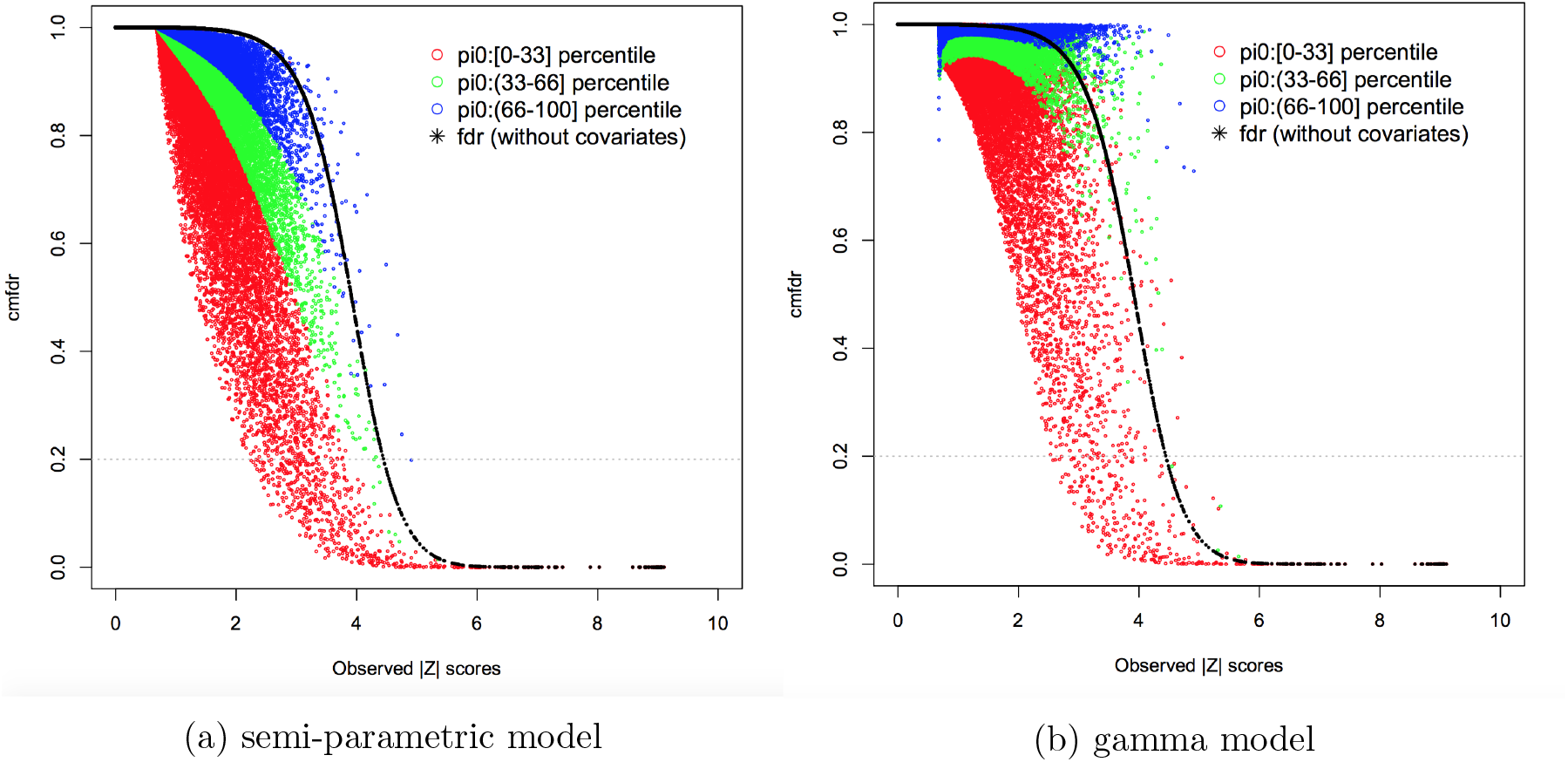
Cmfdr and fdr plotted against observed absolute *z*-scores.

Finally, we compare the non-null densities of semi-parametric (Figure 4a) and gamma (Figure 4b) covariate-modulated mixture models with different values of covariates. The model without covariates is also included (solid lines). Both figures show the non-null densities where all the covariates were set at their corresponding 33 (dash line), 66 (dot line) and 99 (dash-dot line) percentiles. With increasing values for the covariates, the densities show progressively heavier tails. The non-null density of the model without covariates shifts to the right, as compared to the semi-parametric model with covariates in Figure 4a. This shift is probably due to the fact that the variance of the null density (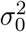) is larger in the model without covariates (median: 1.31, 95% credible interval: 1.29 – 1.33) than the model with covariates (median: 1.12, 95% credible interval: 1.09 – 1.15). The shift also appears in Figure 4b where the median of 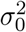 from the gamma model is 1.24 (95% credible interval: 1.22 – 1.26). These results collectively indicate that the enrichment annotation categories we employ here (TotLD, H, and ProteinCoding) provide useful information for selecting “interesting” subsets of SNPs for further analysis.

**Figure 4.**
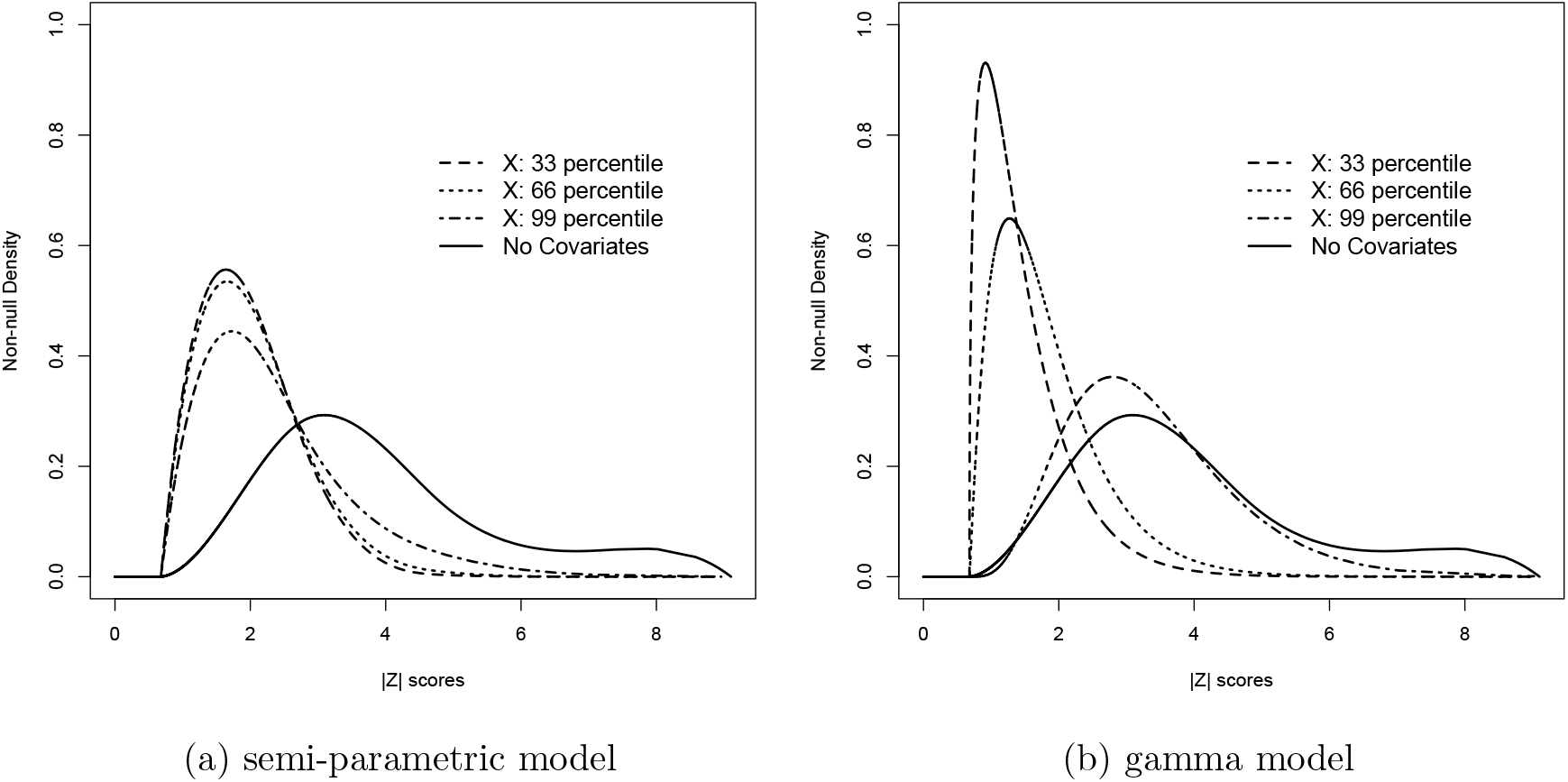
Non-null densities where all three covariates were set at their corresponding 33, 66 and 99 percentiles.

To examine the biological significance of the SNPs, we performed pathway analyses on the 242 gene sets in the KEGG *homo sapiens* pathways database (http://www.kegg.jp/). To perform these pathway analyses, we implemented the ALIGATOR (Holmans et al., 2009) algorithm, which tests for overrepresentation of biological pathways in SNP lists. ALIGATOR corrects for LD between SNPs, variable gene size, and multiple testing of nonindependent pathways. Using the 175 SNPs with fdr ≤ 0.20 results in no pathways with *p*-value ≤ 0.05 (corrected for multiple testing). On the other hand, there were 10 pathways with *p*-values ≤ 0.05 using 588 SNPs with semi-parametric cmfdr ≤ 0.20 (Table 2). The *p*-values using 368 SNPs with gamma cmfdr are also listed for comparison. Axon Guidance is ranked highest in both cmfdr models. The 10 top ranked pathways from semi-parametric cmfdr given in Table 2 provide interesting insight into the pathogenesis of schizophrenia, given that the KEGG database is expertly curated without prior emphasis in terms of disease etiology. The top ranked pathways show abnormal axonal connectivity, lipid metabolizing, and voltage-gated ion channels, as well as comorbid conditions that have been noted among patients with schizophrenia in prior research (Greiner and Nicolson, 1965; Lidow, 2003; Battaglino et al., 2004; Leucht et al., 2007; Putnam, Sun and Zhao, 2011; Maiti et al., 2011; Buckley, Pillai and Howell, 2011; Gardiner et al., 2012; Liu et al., 2013). A complete list of the 242 KEGG *homo sapien* pathways and their ALIGATOR *p*-values are given in Supplementary 2.

**Table 2.**
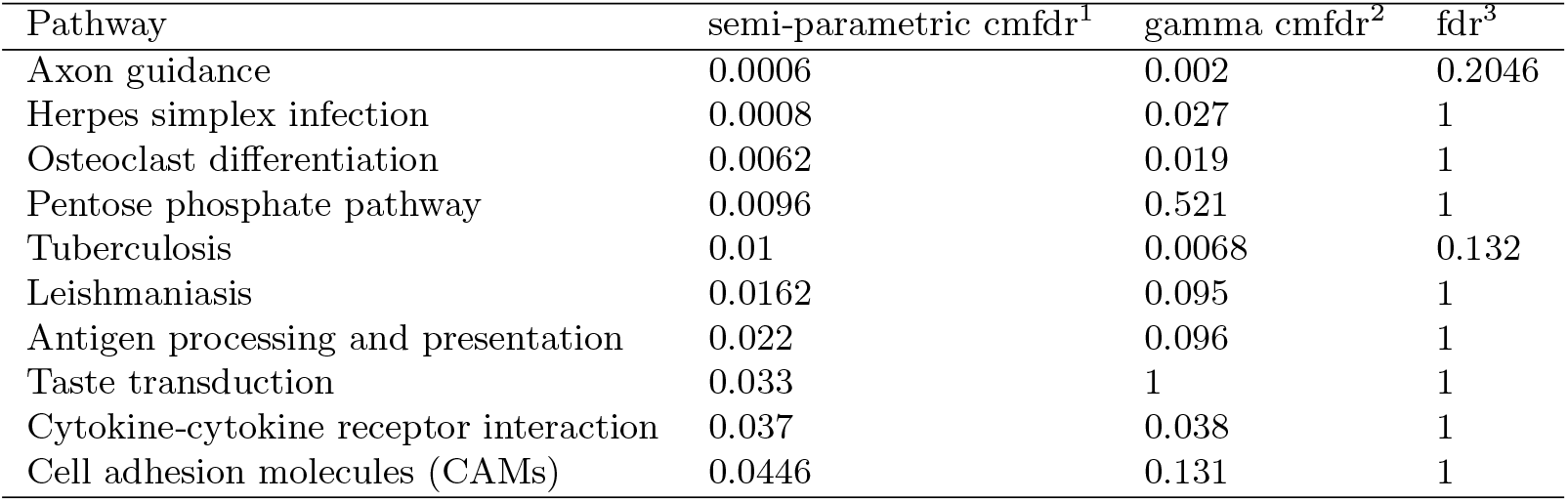
KEGG PATHWAY with ALIGATOR *p*-values from three models

## 4 Discussion

GWAS of highly polygenic traits such as schizophrenia remain underpowered to detect most genetic variants involved in the disorder, even with very large sample sizes. By incorporating auxiliary information, the process of gene discovery can be sped up significantly, along with the assessment of the role of molecular pathways. Moreover, the examination of which auxiliary information is useful for predicting non-null status can be informative of the genetic architecture of polygenic traits.

Using a set of genetic loci (SNPs) pruned for approximate independence, we demonstrate a large increase in power in the PGC schizophrenia data using our semi-parametric cmfdr model compared with fdr, as well as previous models for cmfdr that either use a parametric model (gamma fdr) or a model that does not incorporate covariate effects (FDRreg) into the estimation of the non-null density. For example, using a 0.20 cut-off, we reject 588 null hypotheses with cmfdr compared with only 175 using fdr, or over 3.4 times as many SNPs as the intercept only model, with a similarly large increase in power vs. FDRreg, and a smaller but still substantial increase in power over gamma fdr. This increase in power appears to be driven by a better-fitting model of the tails of the non-null distribution for highly enriched SNPs.

Our choice of covariates in the PGC schizophrenia application was driven by scientific considerations based on theory and substantial prior evidence that these annotations enriched for non-null associations (Schork et al., 2013). In general, we recommend selection of covariates based on these criteria. However, the model could also be used for exploratory analyses, to examine whether a given annotation significantly enriches for associations. For this use, it would be useful to implement a model-selection metric such as the Watanabe-Akaike Information Criterion (WAIC, Vehtari and Gelman (2014)).

The proposed cmfdr model assumes independence of the *z*-scores. To ensure this was approximately true in the current data example, we randomly pruned SNPs so that no two SNPs in the sample were correlated at more than *r*^2^ = 0.20. We thus need to delete many tests to achieve independence. Our current research considers alternative schemes to explicitly model the effects of the correlation on the values of the *z*-scores. We are also developing an extension of the cmfdr model that also incorporates biological networks (gene sets with graphical model structure determined by biological interactions).

## Acknowledgements

The authors are grateful to the anonymous Associate Editor and two referees for their insightful reviews and comments, which greatly improved the paper. The study is supported by NIH grant R01GM104400.

## SUPPLEMENTARY MATERIAL

**Supplementary materials for “Semi-parametric covariate-modulated local false discovery rate for genome-wide association studies”:**

(). The supplement consists of 4 sections. Section 1 presents conditional posteriors and Gibbs sampling algorithm. Section 2 provides the full list of KEGG *homo sapiens* pathways with ALIGATOR *p*-values from different models. Section 3 demonstrates identifiability of the mixture model. Section 4 shows convergence diagnosis plots of parameter estimates.

